# A cyclin-dependent kinase 5-derived peptide inhibits Cdk5/p25 activity and improves neurodegenerative phenotypes

**DOI:** 10.1101/2020.05.12.090472

**Authors:** Jinsoo Seo, Ping-Chieh Pao, Oleg Kritskiy, Audrey Lee, Debasis Patnaik, L. Ashely Watson, Michael Bula, Scarlett J. Barker, Jay Penney, M. Catarina Silva, Stephen J. Haggarty, Li-Huei Tsai

## Abstract

Aberrant activity of cyclin-dependent kinase (Cdk5) has been implicated in various neurodegenerative diseases. This effect is mediated by pathological cleavage of the Cdk5 activator p35 to produce the truncated product p25, exhibiting increased stability and altered substrate specificity. The benefit of blocking p25 production has been demonstrated in various rodent and human neurodegenerative models. However, important Cdk5/p35 functions in the developing and adult brain have made it challenging to selectively target the detrimental effects of Cdk5/p25 while sparing the physiological functions of Cdk5/p35. Here, we report a 12-amino acid-long peptide fragment derived from Cdk5 (the Cdk5 inhibitory (Cdk5i) peptide) that shows a high binding affinity toward the Cdk5/p25 complex and can efficiently and selectively inhibit Cdk5/p25 kinase activity. Using cellular assays, mouse neurodegeneration models and human cerebral organoids generated from patient-derived iPSCs, we demonstrate beneficial effects of the Cdk5i peptide on various pathological phenotypes including gliosis, DNA damage, and Tau hyperphosphorylation.

## Introduction

Unlike other cyclin-dependent kinases (Cdks), Cdk5 is highly expressed in neurons along with its activators p35 and p39. This proline-directed serine/threonine kinase is involved in various signaling pathways in the brain, regulating cortical development, axonal guidance, synaptic plasticity, and learning and memory (Chae et al., 1997; Gilmore et al., 1998; Kim and Ryan, 2010; Seo et al., 2014). The critical importance of Cdk5 for brain development and function was revealed by the studies using transgenic mice lacking its expression; Cdk5 KO mice display lethality at embryonic stages with neuronal migration defects in the cortex and hippocampus among other brain areas (Ohshima et al., 1996; Gilmore et al., 1998). These observations suggested not only a pivotal role for Cdk5 in brain development, but also that its inhibition might cause detrimental effects.

Dysregulation of Cdk5 activity has been associated with multiple types of brain pathology including neurodegenerative diseases such as Alzheimer’s disease and Parkinson’s disease (Patrick et al., 1999; Lee et al., 2000; Smith et al., 2003; Seo et al., 2014; Park et al., 2019). One prominent avenue leading to Cdk5 dysregulation is via cleavage of the Cdk5 activator p35 to p25. Various neurodegenerative stimuli, including protein aggregates and oxidative stress can trigger abnormal increases in intracellular calcium levels leading to activation of the calcium-dependent cysteine protease, calpain. Calpain in turn acts on p35, cleaving its N-terminal regulatory motif and generating elevated levels of the p25 proteolytic fragment. p25 maintains the ability to activate Cdk5 but with a longer half-life compared to Cdk5/p35. Calpain-mediated cleavage also removes myristoylated residues that normally tether Cdk5/p35 to membrane sites, liberating Cdk5 and allowing the Cdk5/p25 complex to access additional kinase substrates (Kusakawa et al., 2000; Lee et al., 2000; Su and Tsai, 2011).

Lack of auto-inhibition and distinct subcellular localization of Cdk5/p25 compared to Cdk5/p35 suggest the need for a tailored approach to specifically target Cdk5/p25 for Cdk5-mediated pathology. However, because p25 is a derivative of p35, it is challenging to design inhibitors targeting Cdk5/p25 without affecting Cdk5/p35. Genetic manipulation to generate mutant p35 that is non-cleavable by calpain but still functional (Δp35) has demonstrated the efficacy of blocking p25 activity (Seo et al., 2014). Transgenic mice in which endogenous p35 was replaced with Δp35 were shown to be less susceptible to Alzheimer’s disease-related pathology and tauopathy when they were crossed with 5XFAD mice (expressing five familial AD-linked mutations - *APP* KM670/671NL, *APP* I716V, *APP* V717I, *PSEN1* M146L and *PSEN1* L286V) and Tau P301S mice (expressing *MAPT* carrying the P301S mutation) (Seo et al., 2014; 2017). While the Δp35 transgenic model, effectively a p25 knockout, has been critical to understanding the physiological and pathological functions of Cdk5/p25, the translational potential of this approach is very limited. Small molecules such as roscovitine, a kinase inhibitor with specificity for multiple Cdk family members, have also been utilized to study Cdk5 functions. The utility of roscovitine is limited, however, by a lack of specificity for Cdk5 relative to other Cdk family members (including Cdk1, Cdk2, Cdk7 and Cdk9) (Cicenas et al., 2015).

An alternative approach has been the design of peptides to inhibit Cdk5/p25 function. Multiple truncated peptides derived from p35, including CIP (Cdk5 inhibitory peptide) and P5, have been tested, in some cases showing inhibition of Cdk5 activity (Zheng et al., 2005; 2010). Among them, the shortest peptide (P5; 24 amino-acids), showed the most effectiveness. A modified form of P5, with C-terminal cell-penetrating transactivator of transcription (TAT) sequences and an N-terminal FITC tag, was further utilized for *in vivo* investigation in several neurodegenerative mouse models. Injection of this modified P5 into 5XFAD mice and CK-p25 mice (overexpressing p25 in forebrain neurons), reduced the hyperactivation of Cdk5 and gliosis, and restored synaptic and cognitive functions (Shukla et al., 2013; 2017).

Such a peptide-based approach comes with many advantages. As applied in the studies mentioned above, peptides with some modifications can be directly administrated to rodent brains to monitor their actions. Fluorescence polarization assays can likewise be designed to identify small molecule inhibitors mimicking the effects of the peptide in a high-throughput manner (Moerke, 2009; Wei et al., 2015; Hall et al., 2016). It is worth noting that the length of a peptide is the limiting factor in such assays; shorter peptide lengths increased assay efficiency.

Our earlier efforts determined the structural basis of the Cdk5/p25 interaction (Tarricone et al., 2001) and provided insight for developing a novel inhibitory peptide for Cdk5/p25 derived from the Cdk5 sequence which we report here. This 12-amino-acid long Cdk5 inhibitory peptide (hereafter referred to as ‘Cdk5i peptide’) has higher binding affinity for Cdk5/p25 than Cdk5 alone or Cdk2, and displays Cdk5/p25-specific inhibitory function without affecting physiological activities of Cdk5 or Cdk2. Importantly, when delivered to mouse models of neurodegenerative disease (Tau P301S and CK-p25 mice), and human cerebral organoids generated from iPSC of patient with frontotemporal dementia (FTD) carrying the *MAPT* P301L mutation, the Cdk5i peptide ameliorated multiple pathological phenotypes associated with neurodegeneration.

## Materials & Methods

### Cdk5i and scrambled peptides

Cdk5i peptide consists of 12 amino acids (ARAFGIPVRCYS) that is derived from a activation loop (T-loop) of Cdk5 known as the important site for interacting p25 (Tarricone et al., 2001). The sequence for the scrambled is AFRSPCARIGYV. For pull-down assay, biotin was tagged at N-terminus of both Cdk5i and scrambled peptides. For intraperitoneal (I.P.) injection to mice and treatment to iPSCs-derived cerebral organoids, the peptides were conjugated with FITC along with a linker (Ahx-GGG) at N-terminus and the TAT-sequence (YGRKKRRQRRR) was tagged at C-terminus. All peptides were synthesized by Peptide 2.0 Inc (Chantilly, VA).

### Recombinant p25/Cdk5 for Cdk5 kinase activity assays

Recombinant human Cdk5 and p25 with N-terminal GST-tags were expressed by Sf9 insect cells via a baculovirus expression system followed by GST affinity purification (BPS Bioscience, San Diego CA).

### Microscale thermophoresis-based biophysical assays

The binding affinity of Cdk5i peptide was determined by microscale thermophoresis (MST) using reagents, consumables, and Monolith NT.115Pico from NanoTemper Technologies (Munich, Germany). The recombinant CDK5/p25 protein prep with a HIS was kindly provided by Dr. Kenneth S. Kosik, UC Santa Barbara. 6X-HIS tagged CDK5 (CDK5-236H), and 6X-HIS tagged Cdk2 (CDK2-655H) were purchased from Creative Biomart (https://www.creativebiomart.net/). The HIS tagged protein samples were labeled with the red fluorescent dye NT647 using the Monolith NT HIS-tag labeling kit RED-tris-NTA. A serial dilution of the small molecule was made at 2X concentration with the MST assay buffer (50 mM Tris pH 7.4, 150 mM NaCl, 10 mM MgCl_2_ and 0.01% Brij-35) and subsequently mixed with an equal volume of 2X concentration of the labeled fluorescent protein. The labeled protein and the Cdk5i peptide mix were allowed to incubate, before filling the capillaries. MST assays were performed at a final concentration of 10 nM labeled protein, with 20% LED/excitation power and high MST power (60%) using regular capillaries for Monolith NT.115. Premium capillaries were used to perform MST with the labeled CDK5/p25 prep to improve the quality of data. The MST results were evaluated with the NanoTemper’s MO Affinity Analysis software to estimate the *K*_d._

### Biolayer interferometry-based biophysical assays

The binding interaction between the Cdk5i peptide with the recombinant Cdk5/p25 protein was also investigated via biolayer interferometry (BLI) using the Octet Red 384 instrument (ForteBio, Fremont, California) with 1X PBS with 0.01% Brij-35 as the assay buffer. The same recombinant Cdk5/p25 protein used for the MST assays was labeled with biotin using EZ-LinkNHS-PEG4-Biotinylation Kit (Thermo Scientific, Waltham, MA) and excess reagent was removed with a spin desalting column as per the manufacturer’s instruction. The biotinylated recombinant Cdk5/p25 protein for BLI was eluted in 1X phosphate buffered saline (PBS). Streptavidin (SA) sensors were utilized to study the biophysical interaction between the Cdk5i peptide and biotinylated Cdk5/p25 via BLI. Before starting the BLI experiment, the streptavidin sensors were soaked by dipping in 200 μl of assay buffer in a 96-well Greiner Bio-One Black flat bottom plate (#655209). BLI analysis was performed in 80 μL volume in Greiner Bio-One 384-well black flat bottom PP plates (#781209, Greiner, Monroe, North Carolina) with an initial baseline step, followed by loading of 250 nM biotinylated Cdk5/p25. The recombinant protein and the sample analyte (n=5) were organized in a 384-well plate as per a plate map recommended for the 8-channel mode kinetic analysis, where the streptavidin sensors step from low to high concentration of the Cdk5i sample analyte. Subsequent steps included a second baseline (120s), association (240s), and dissociation (240s) for the analysis of samples. All the sensors were loaded with the recombinant biotinylated Cdk5/p25 protein, and three sensors with the comparable concentration of DMSO in assay buffer was used as the reference (n=3). The results were analyzed with the Data Acquisition HT 11.0 (Fortebio) following reference subtraction (an average of three sensors) with the 1:1 binding model and fitted globally to estimate the equilibrium dissociation constant.

### Pull-down assay

Recombinant Cdk5/p25 complex or whole brain lysates were incubated with biotinylated Cdk5i or scrambled peptide in binding buffer (50 mM Tris, 150 mM NaCl, 0.1% NP-40, pH 7.5) for overnight at 4°C. The following day, Streptavidin-coated beads (GE Life science) were prepared and washed with binding buffer three times. After the final wash, beads were resuspended to produce a 50% slurry by volume, and added to recombinant Cdk5/p25 complex. After rotating the mixture at 4°C for 1 hour, samples were centrifuged at 2,500 rpm for 2 min, and the supernatant were saved as ‘flow through’. Resuspended beads were subjected for immunoblotting to detect the interaction between the peptides and Cdk5, Cdk2, p25 or p35.

### IP-linked kinase assay

Recombinant Cdk5/p25 complex or whole mouse brain lysates were incubated with anti-Cdk5 or anti-Cdk2 antibodies for overnight at 4°C and the immunocomplex was prepared as described previously (Seo et al., 2014), and loaded on SDS-PAGE gel and resolved immediately. After running, the gel was carefully dried with plastic wrap and filter paper to avoid wrinkles, and the dry gel was exposed onto a phosphoscreen overnight. The next day, amount of ^32^P-Histone H1 was measured using a phosphoimager (GE Typhoon). Coomassie staining was performed to confirm that equal amount of proteins was loaded for each group.

### Animals

All animal experiments were performed with approval from the MIT Committee on Animal Care (CAC). CK-p25 mice were raised on doxycycline containing diet until 2 months of age and then switched to normal diet for two weeks along with I.P. injection of Cdk5i or scrambled peptide. Tau P301S (PS19 line) (Yoshiyama et al., 2007), and wild-type mice were obtained from the Jackson Laboratory (Bar Harbor, ME).

### Primary Neuronal Culture

Primary neurons were generated from the cortical region of timed pregnancy mice (Swiss-Webster) on day E16. Cells were dissociated into a single cell suspension and maintained in Neurobasal Media (Thermo Fisher Scientific) supplemented with L-glutamine (5 mM), penicillin and streptomycin, and B27 neuronal additive. Cells were plated at 2.5×10^6^ cells/10-cm dish, and then transduced with human 4R0N Tau carrying a P301L mutation at 5 days *in vitro* (DIV). Cultured neurons were treated with Cdk5i or scrambled peptide at a concentration of 10 nM for 6 hours on 14 DIV.

### iPSCs cultures

FTD patient (*MAPT* P301L +/-)-derived iPSC line (Seo et al., 2017) and its isogenic line in which P301L mutation in *MAPT* was corrected by CRISPR/Cas9 genome editing were cultured on irradiated mouse embryonic fibroblasts (MEFs) in DMEM/F12, HEPES media (Gibco) supplemented with 20% knockout serum replacement (KSR) (Gibco), 1X non-essential amino acids (NEAA), 1X GlutaMAX, (Life Technologies) β-fibroblast growth factor (FGF2, PeproTech) and 0.1 mM 2-mercaptoethanol (Sigma-Aldrich). Original approval for human subject work was obtained under a Partners/Massachusetts General Hospital-approved Institutional Review Board Protocol (#2010P001611/MGH).

### Generation of isogenic control line

A sgRNA for targeting the P301L site in *MAPT* gene (AACATTAAGCATGTCCCGGGAGG) was designed by http://crispr.mit.edu. And both sgRNA and repair template were synthesized by IDT (Integrated DNA Technologies). The CRISPR/Cas9 plasmid (pSpCas9-2A-GFP, PX458) was purchased from Addgene and the sgRNA was cloned into the plasmid and electroporated into iPSCs followed by FACS and colony inspection as described previously (Ran et al., 2013; Seo et al., 2017).

### Cerebral organoid culture

Cerebral organoids were generated from iPSCs as described previously (Raja et al., 2016; Seo et al., 2017). In brief, embryoid bodies (EBs) were formed by loading 12,000 iPSCs per well into 96-well plates pre-coated with Pluronic acid (1%, F-127, Sigma-Aldrich) and maintained in media consists of Glasgow-MEM supplement with 20% KSR, 1X sodium pyruvate, 1X NEAA, 0.1 mM 2-mercaptoethanol, 20 μM ROCK inhibitor (Y-27632, Millipore), 5 μM TGFβ-inhibitor (SB431532, Tocris Biosciences), 3 μM Wnt-inhibitor (IWRe1, Tocris Biosciences) for 20 days. Cerebral organoids were then transferred to non-adherent petri-dishes and cultured in media consisted of DMEM/F12 supplemented with 1X Chemically Defined Lipid Concentrate and 1X N2-supplement in 40% O_2_/5% CO_2_ conditions to promote neuroepithelial formation. From day 35, 5 μM heparin (Sigma-Aldrich), 10% FBS and 1% matrigel (Life Sciences) were added to the medium.

### Immunoblot Analysis

Cultured neurons or homogenized brain lysates were washed once with cold 1 X PBS and lysed in RIPA buffer (150 mM NaCl, 1% IGEPAL, 0.5% sodium deoxycholate, 0.1% SDS, and 50 mM Tris-HCl, pH 8.0, supplemented with protease and phosphatase inhibitors). Supernatant was collected after centrifugation at 12,000 x g for 10 min at 4 °C. Samples were loaded in equal amounts of protein onto a 10% polyacrylamide gel. After electrophoresis, proteins were transferred to Immobilon PVDF 0.45 μm membranes (Millipore). Nonspecific binding was blocked by 5% Bovine Serum Albumin in Tris-buffered saline, 0.05% Tween 20. Membranes were incubated with primary antibody for overnight at 4 °C, and secondary antibodies for 1 hour at RT. Band intensity was analyzed using ImageJ software.

### Immunohistochemistry

Mice were transcardially perfused with 4% paraformaldehyde in PBS under anesthesia (2:1 of ketamine/xylazine), and the brains were sectioned at 40 μm thickness with a Leica VT1000S vibratome (Leica). Cerebral organoids were fixed with 4% paraformaldehyde for overnight, and transferred to 30% sucrose in PBS and kept until they sink. Dehydrated cerebral organoids were cryosectioned at 40 μm thickness. Sections were permeabilized and blocked in PBS containing 0.2% TritonX-100 and 10% normal donkey serum for 1 hour at RT, then incubated with primary antibody at 4°C overnight. Primary antibody was visualized using the appropriate secondary antibody conjugated to Alexa Fluor 488, Alexa Fluor 594, or Alexa Fluor 647 (Thermo Fisher Scientific). Nuclei were visualized with Hoechst 33342 (Thermo Fisher Scientific).

### Microscopy

All images were captured using a Zeiss LSM 710 confocal microscope and the ZEN software, and analyzed using the ImageJ software (National Institutes of Health, https://imagej.nih.gov/ij/, RRID: SCR_003070).

### Tau seeding activity assay

Whole brain lysates from Tau P301S mice with Cdk5i or scrambled treated and FRET biosensor cell lines described previously were prepared (Seo et al., 2017). For the assay, cells were plated in a 96-well plate at a density of 40,000 cells/well. Brain homogenate samples were transduced into cells 16 hours later plating at 50% confluence, using 1 μL Lipofectamine/well. After 48 hours incubation, cells were harvested and fixed in 4% paraformaldehyde. An LSR II HST-2 flow cytometer was used to measure the FRET signal in each sample.

### Antibodies

Cdk5 (C-8, Santa Cruz Biotechnology), p35 (Tsai lab), Cdk2 (SC-163, Santa Cruz Biotechnology), FITC (71-1900, Thermo Fischer Scientific), γ-H2AX (05-636, Millipore), NeuN (266 004, Synaptic Systems), pTau T181 (9632, Cell Signaling Technology), pTau S396 (5383, Cell Signaling Technology), pTau S404 (35834, Cell Signaling Technology), GAPDH (2118, Cell Signaling Technology).

## Results

### Design of Cdk5/p25 complex-specific inhibitory peptide

To develop an inhibitory peptide specific to the Cdk5/p25 complex, we utilized our previous work elucidating the crystal structure of the Cdk5/p25 complex, which revealed a critical region in Cdk5 required for the interaction with p25 (Tarricone et al., 2001). This study found by cross-species comparison that the activation loop (T-loop) of Cdk5 is highly conserved from mammals to insects (human, mouse, *Xenopus, C. elegans, Drosophila*), but distinct from that of other cyclin-dependent kinases such as Cdk2. We thus selected a 12-amino acid sequence (ARAFGIPVRCYS) from the Cdk5 T-loop to generate the Cdk5i peptide (**Figure 1A**). The Cdk5i sequence includes residues R149, A150, I153, C157 and S159, all directed toward the p25 surface and predicted to mediate intermolecular interactions between Cdk5 and p25 (**Figure 1A**).

**Figure 1.**
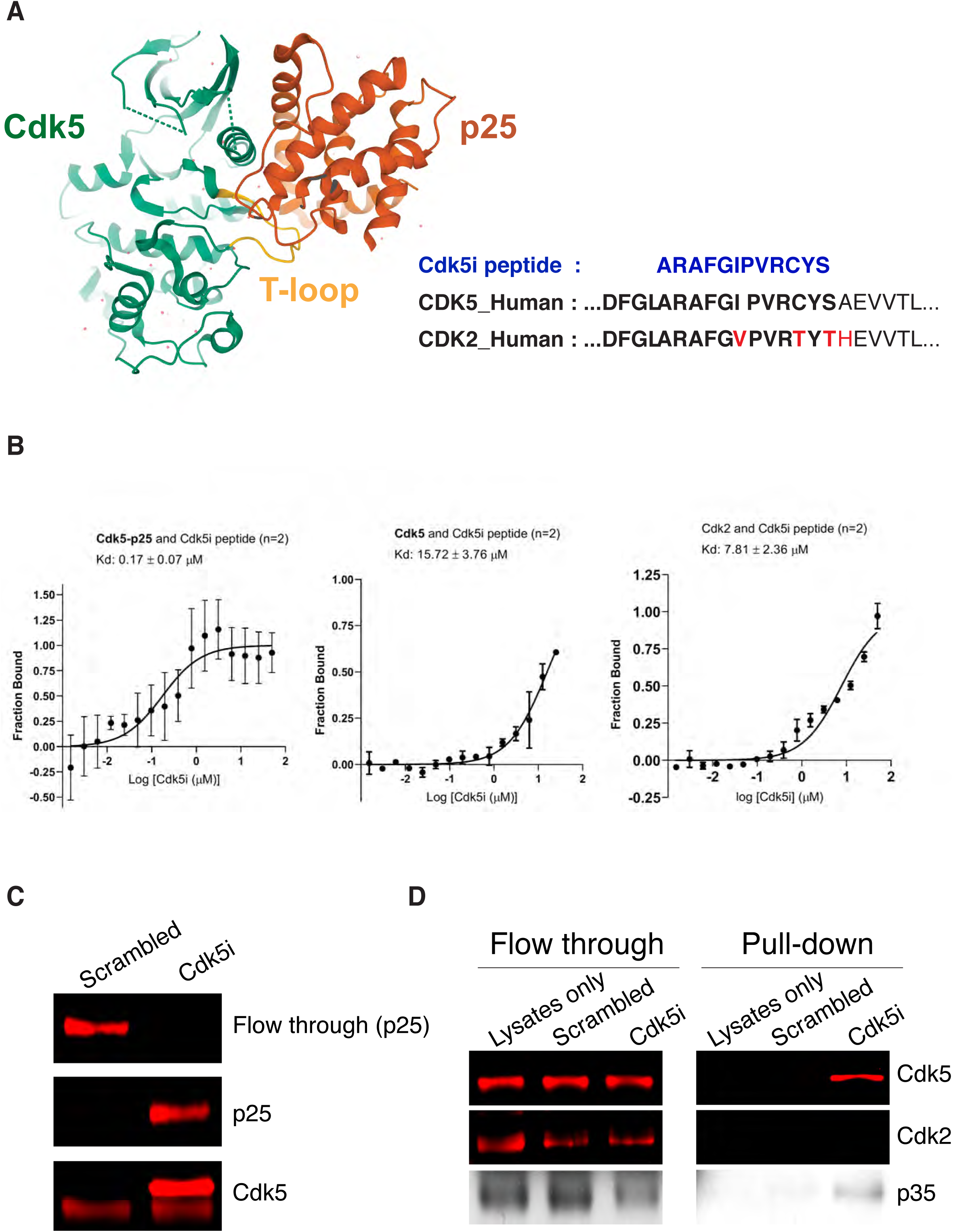
Design of Cdk5/p25 complex-specific inhibitory peptide. **A**, Strategy for the design of Cdk5/p25 complex-specific inhibitory peptide (Cdk5i peptide). The structure of Cdk5/p25 complex was downloaded from RCSB protein data bank. **B**, Microscale thermophoresis-based biophysical assays assessing binding of Cdk5i peptide to recombinant proteins. **C**, Recombinant Cdk5/p25 complex was incubated with biotin-conjugated Cdk5i or scrambled peptide for overnight at 4°C, and subjected to pull-down assay with streptavidin-coated beads. Interaction between the peptides and Cdk5, or p25 was addressed by immunoblotting. **D**, Brain lysates from wild-type mice were incubated with Cdk5i peptide or scrambled for overnight at 4°C, and subjected to pull-down assay. Interaction between the peptides and Cdk5, Cdk2 or p35 was addressed by immunoblotting.

To begin to characterize the strength and specificity of Cdk5i peptide binding, we first utilized an *in vitro* binding assay based upon microscale thermophoresis to probe interactions of the peptide with recombinant Cdk5, Cdk2 or Cdk5/p25 complex. We found that the Cdk5i peptide bound to the Cdk5/p25 complex with a K_d_ = 0.17 μM, which was more than 40-fold more strongly to either Cdk5 or Cdk2 alone (**Figure 1B**). As an independent biophysical assay, we also verified the binding interaction between the Cdk5i peptide and recombinant Cdk5/p25 protein using biolayer interferometry yielding an affinity measure (K_d_ = 0.22 μM, **Supplemental Figure 1**). We next utilized pull-down assays using recombinant proteins or brain lysate to probe Cdk5i peptide binding. As a control for peptide sequence specificity, we also included a scrambled peptide (AFRSPCARIGYV) in these experiments. First, the Cdk5i or scrambled peptides were incubated with recombinant Cdk5/p25 complex followed by pull-down. As a bait, the Cdk5i and scrambled peptides were tagged with biotin at their N-terminal and captured by streptavidin-coated beads. We found that the Cdk5i peptide pulled down both Cdk5 and p25 while the scrambled peptide exhibited minimal binding to either protein (**Figure 1C**).

We then subjected wild-type mouse brain lysates to similar pull-down with the Cdk5i or scrambled peptides to assay their interaction with Cdk5, Cdk2, p25 and p35. We detected no interaction between the scrambled peptide and Cdk5 or Cdk2, while the Cdk5i peptide specifically pulled down Cdk5 but not Cdk2 (**Figure 1D**). p25 and p35 can be detected as separate sized bands on western blot with our p35 antibody, however, p25 levels are generally very low under normal physiological conditions. Consistently, we were unable to detect p25 in our brain lysate or following pull-down with the Cdk5i or scrambled peptides (**Figure 1D**). In contrast, we found that p35 exhibited interaction with the Cdk5i but not the scrambled peptide under these conditions (**Figure 1D**). Together, these findings show that the Cdk5i peptide binds readily to both recombinant and brain-derived Cdk5 protein and exhibits strong selectivity for Cdk5 over Cdk2.

### Inhibitory action of the Cdk5i peptide on Cdk5/p25 complex function

Our *in vitro* binding and pull-down assays demonstrate strong affinity of our Cdk5i peptide for Cdk5 and Cdk5/p25, thus we next wanted to assess whether the Cdk5i peptide exerts an effect on the kinase activity of Cdk5 complexes. We first examined the effect of Cdk5i peptide on kinase activity of the Cdk5/p25 complex prepared from recombinant systems. Compared to Cdk5/p25 complex incubated with scrambled peptide, Cdk5/p25 incubated with Cdk5i peptide showed significantly reduced ability to phosphorylate histone H1, a Cdk substrate (**Figure 2A**).

**Figure 2.**
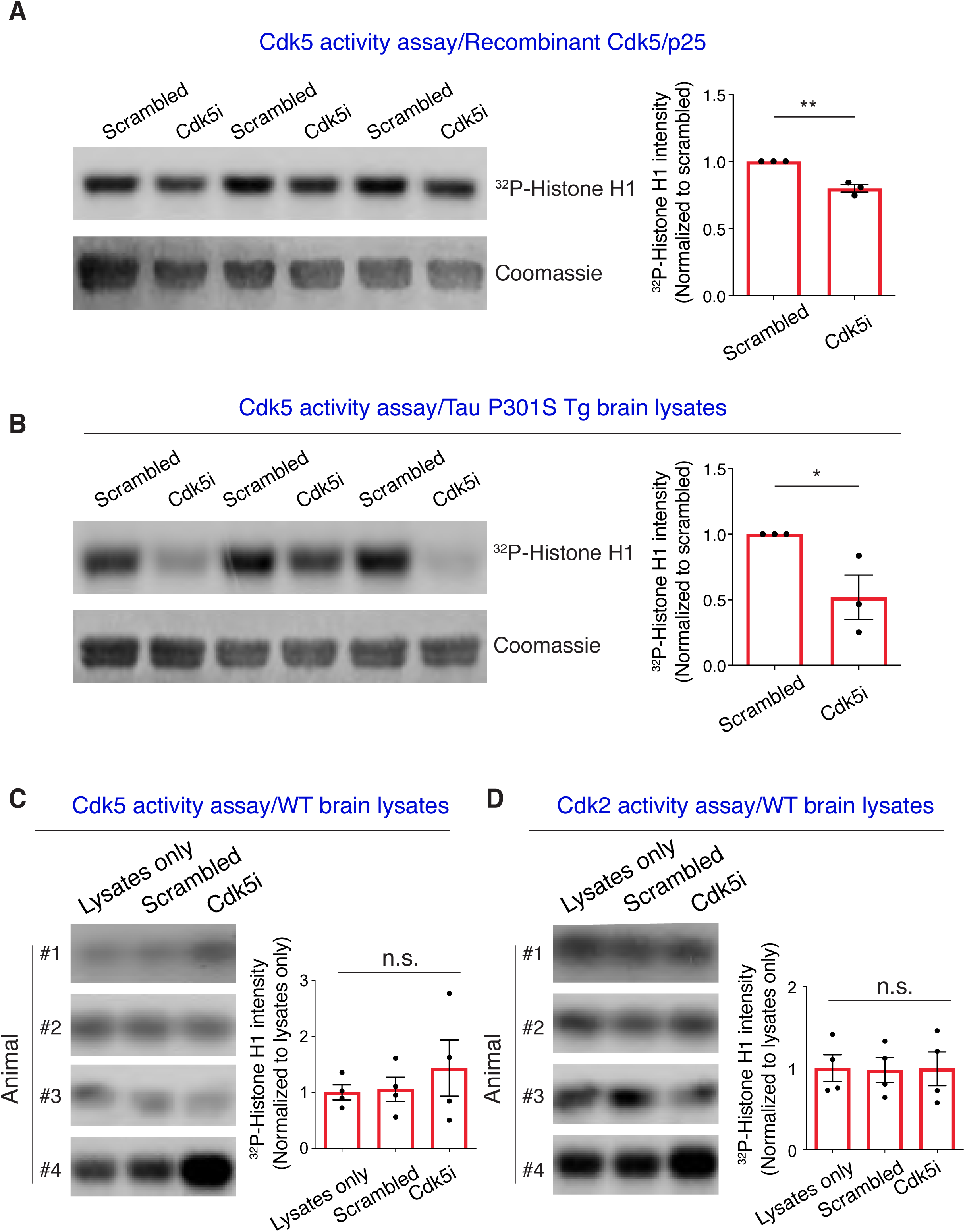
Inhibitory action of Cdk5i peptide on Cdk5/p25 complex. **A**, Recombinant Cdk5/p25 complex was incubated with Cdk5i peptide or scrambled overnight at 4°C, and subjected to IP-linked Cdk5 kinase assay. The bar graph represents the relative radioactivity of ^32^P incorporated into histone H1. **B**, Brain lysates from three-month-old Tau P301S mice were incubated with Cdk5i peptide or scrambled overnight at 4°C, and subjected to IP-linked Cdk5 kinase assay. **C-D**, Brain lysates from three-month-old wild-type mice were incubated with Cdk5i peptide or scrambled overnight at 4°C, and subjected to IP-linked kinase assay for Cdk5 or Cdk2.

To test whether the Cdk5i peptide could likewise exert an inhibitory effect on Cdk5 complexes isolated from neurodegenerating mouse brains, we next incubated lysates from Tau P301S mice with Cdk5i or scrambled peptide and performed a Cdk5 kinase assay. We previously reported that inhibition of Cdk5/p25 activity by replacing endogenous p35 with the non-cleavable Δp35 in this mouse model reduced Cdk5 activity by about 60% (Seo et al., 2017). Consistently, we found a roughly 50% reduction of Cdk5 activity in P301S mice brain lysates after treatment with the Cdk5i peptide compared to those treated with scrambled peptide (**Figure 2B**).

To examine whether the Cdk5i peptide can inhibit Cdk5/p35 function, which is known to be critical for neurodevelopment and synaptic function under physiological conditions, we collected brain lysates from wild-type mice. Following incubation of these lysates with either Cdk5i or scrambled peptides, we measured kinase activity for Cdk5. Despite showing physical interaction with p35 from mouse brain (**Figure 1D**), we found that neither the Cdk5i nor scrambled peptides altered basal Cdk5 kinase activity in wild-type brain lysates (**Figure 2C**). Furthermore, Cdk2 activity in wild-type brain lysates was also unaffected by Cdk5i or scrambled peptides (**Figure 2D**). Thus, the Cdk5i peptide can inhibit the kinase activity of recombinant Cdk5/p25, as well as reducing Cdk5-mediated phosphorylation in Tau P301S but not wild-type brain lysates.

### Preventive effects of the Cdk5i peptide on pathological phenotypes in CK-p25 mice

Multiple pathological effects of Cdk5/p25 signaling, as well as the beneficial effects of inhibiting Cdk5/p25 function have been reported in multiple neurodegenerative conditions including Alzheimer’s disease and Frontotemporal dementia (FTD) (Shukla et al., 2013; Seo et al., 2014; 2017; Shukla et al., 2017). Thus, we sought to examine the effects of our Cdk5i peptide in neurodegeneration-related phenotypes *in vivo*. To that end, we added the TAT sequence (YGRKKRRQRRR) to the C-terminus of the peptide along with conjugating FITC at the N-terminus. The modified version of the peptide was intraperitoneally (IP) injected (40 mg/kg body weight) into CK-p25 mice, an inducible model of AD-like neurodegeneration. We were able to detect FITC signal in both hippocampus and cortex 24 hours after a single injection, indicating successful penetration into the brain (**Figure 3A**).

**Figure 3.**
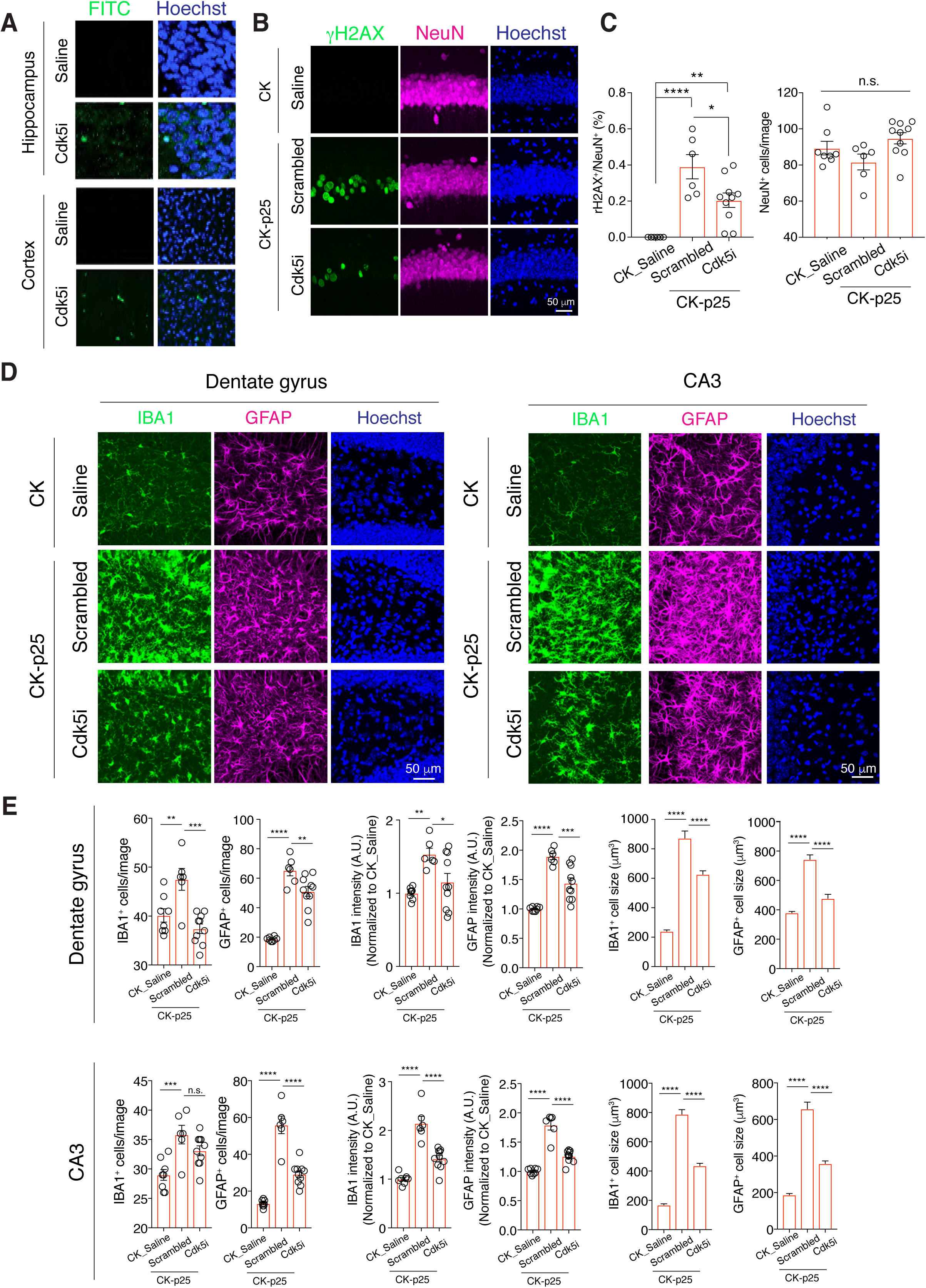
Preventive effects of Cdk5i peptide on pathological phenotypes in CK-p25 mice brain. **A**, FITC-conjugated Cdk5i peptide was injected to three-month-old CK-p25 mice (IP 40 mg/kg body weight) and its expression in hippocampus and cortex was examined using an anti-FITC antibody. **B**, Immunostaining for γ-H2AX and NeuN were performed in hippocampal CA1 region of CK-p25 mice treated with Cdk5i peptide or the scrambled during two weeks of p25 overexpression. **C**, Bar graphs represent the population of γ-H2AX^+^; NeuN^+^ cells out of total NeuN^+^ and the number of NeuN^+^ cells in CA1 of CK-p25 treated with Cdk5i or scrambled peptide. **D**, Immunostaining for GFAP and IBA-1 were performed in hippocampal dentate gyrus region and CA3 of CK-p25 mice treated with Cdk5i peptide or the scrambled during two weeks of p25 overexpression **E**, Bar graphs represent the number of GFAP-, Iba-1-positive cells, levels of GFAP, IBA-1, or size of GFAP-, IBA-1-positive cells in dentate gyrus and CA3 of CK-p25 mice treated with Cdk5i peptide or the scrambled during two weeks of p25 overexpression.

Two weeks of p25 induction in CK-p25 mice has been shown to be sufficient to induce DNA damage, marking the beginning of a cascade leading to neuronal death (Kim et al., 2008). To test whether Cdk5i peptide treatment could counteract p25-induced DNA damage, we treated CK-p25 mice with the Cdk5i or scrambled peptides three times weekly (IP, 40 mg/kg body weight) during the two-week p25 induction period. Consistent with previous reports, we observed prominent accumulation of DNA damage, marked by immunoreactivity for phosphoserine 139 of histone H2AX (γH2AX), in the CA1 area of hippocampus after two weeks induction (Kim et al., 2008) (**Figure 3B-C**). We found that treatment of CK-p25 mice with the Cdk5i peptide significantly reduced γH2AX signal intensity relative to those receiving the scrambled peptide, though DNA damage remained higher than wild-type mice injected with saline (**Figure 3B-C**).

Profound gliosis is also an early pathological response to p25 induction, thus we examined astrocyte and microglia changes in the hippocampi of CK-p25 mice by immunohistochemistry with anti-GFAP and anti-IBA1 antibodies, respectively. We found that two-week treatment with the Cdk5i peptide significantly reduced levels of GFAP and IBA1 in the dentate gyrus region and CA3 of the hippocampus relative to scrambled peptide (**Figure 3D-E**). Likewise, the increased size of astrocytes and microglia we observed in the dentate gyrus and CA3 of CK-p25 mice was largely reversed by Cdk5i peptide injection, suggesting attenuation of p25-induced gliosis. Taken together, these data suggest the Cdk5i peptide exerts protective effects on Cdk5/p25-mediated neurodegenerative phenotypes *in vivo*.

### The Cdk5i peptide ameliorates Tau hyperphosphorylation in human and rodent FTD models

We next examined the effects of the Cdk5i peptide in tauopathy models which have been strongly associated with Cdk5 hyperactivity (Kimura et al., 2014; Seo et al., 2017). We first overexpressed human 4R0N Tau carrying the FTD-associated P301L mutation in mouse primary neurons starting at DIV 5. Then on DIV 14, we treated cells with either the Cdk5i or scrambled peptide (10 nM) for 6 hours before examining Tau phosphorylation levels by western blotting. Treatment with the Cdk5i peptide induced a significant reduction of Tau phosphorylation on both residues T181 and S396, when compared to treatment with the scrambled peptide (**Figure 4A**).

**Figure 4.**
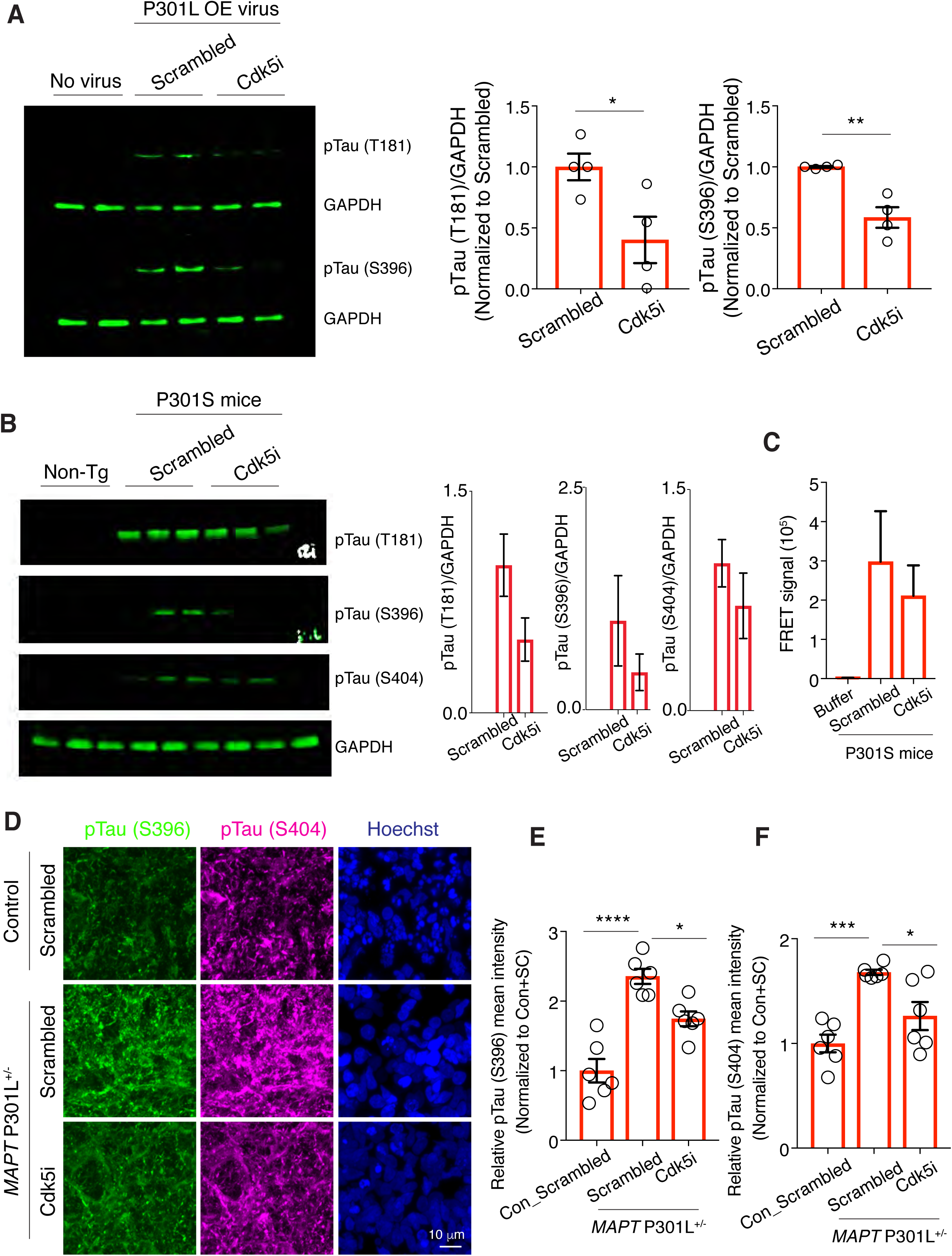
The Cdk5i peptide ameliorates Tau hyperphosphorylation in human and rodent FTD models. **A**, Levels of pTau T181, pTau S396, and GAPDH were measured after Cdk5i or scrambled peptide treatment in mouse primary neurons overexpressing human Tau P301L mutation. The bar graphs represent the quantification of relative immunoreactivity for pTau T181 and pTau S396 normalized to the scrambled-treated group. **B**, Levels of pTau T181, pTau S396, pTau S404, and GAPDH were measured in brain lysates from non-transgenic (lanes 1-3) and Tau P301S mice treated with either scrambled (lanes 4-6) or Cdk5i peptide (lanes 7-9). Three animals in each group were analyzed. The bar graphs represent the quantification of relative immunoreactivity for pTau T181, pTau S396 and pTau S404 normalized to the non-transgenic group. **C**, Four-month-old Tau P301S animals that were treated with either scrambled or Cdk5i peptides (40 mg/kg body weight) for one month followed by cortical tissue harvesting, were used for a Tau seeding assay. Cortical lysates (5 μg) were added to biosensor HEK293T cells, and their Tau seeding activity was assessed by measuring FRET signals. Three animals in each group were analyzed. **D**, Representative images of pTau S396 and pTau S404 in 4.5-month-old cerebral organoids generated from *MAPT* P301L +/- iPSCs with scrambled peptide treatment, *MAPT* P301L +/- iPSCs with Cdk5i peptide treatment and isogenic control iPSCs with scrambled peptide treatment. **E-F**, Bar graphs represent the quantification of immunoreactivity for pTau S396 **(E)** and pTau S404 **(F)** normalized to isogenic control organoids with the scrambled peptide treatment. Three organoids and two sections per organoid were analyzed in each group.

We then examined *in vivo* effects of the Cdk5i peptide using the PS19 mouse model of FTD, overexpressing pathogenic human mutant Tau P301S under the control of the prion promoter (Yoshiyama et al., 2007). These mice display increased levels of phosphorylated Tau in brain, and their brain lysates show increased Tau seeding activity when tested using Tau monoclonal FRET biosensor cells (Holmes et al., 2014). We showed previously that these phenotypes were attenuated when p25 generation is abolished in compound PS19; Δp35 transgenic mice (Seo et al., 2017). Either Cdk5i or scrambled peptides were delivered to 3-month-old PS19 mice three times per week by IP injection for one month. We then performed immunostaining to examine levels of pTau. Consistent with data from our *in vitro* experiments with cultured primary neurons (**Figure 4A**), we observed that levels of pTau at multiple residues (T181, S396 and S404) exhibited strong trends toward reduction by Cdk5i peptide treatment compared to mice injected with the scrambled peptide (**Figure 4B**). Hyperphosphorylation of Tau is known to lead to misfolding and aggregation of Tau by liberating the protein from microtubules and exposing its hydrophobic surface (Mazanetz and Fischer, 2007; Kolarova et al., 2012). This insoluble form of Tau is thought to induce propagation of tauopathy across cells by “seeding” the formation of additional aggregates. Therefore, we next utilized FRET biosensor HEK293T cells expressing human Tau carrying P301S mutation and fused with either CFP or YFP to test Tau seeding activity. Compared to scrambled peptide-treated Tau P301S mouse brains, we observed a trend toward reduced seeding effect from Tau P301S brains treated with the Cdk5i peptide (**Figure 4C**).

We have found striking effects of the Cdk5i peptide treatment in mouse brains and cultured cells. To extend these findings to a human neurodegenerative context, we took advantage of human iPSC models. We utilized iPSCs derived from an FTD patient carrying a heterozygous *MAPT* P301L mutation, and generated isogenic cells where the P301L mutation was corrected by CRISPR/Cas9 genome editing (**Supplementary Figure 2**). We previously reported that cerebral organoids derived from the parental *MAPT* P301L line faithfully recapitulate Tau hyperphosphorylation, as observed in patient brains (Seo et al., 2017). Consistently, in cerebral organoids generated from our isogenic pair of iPSC lines, we found increased levels of pTau (pS396 and pS404) in 4-month-old organoids carrying the *MAPT* P301L compared to corrected controls. Importantly, treatment of *MAPT* P301L cerebral organoids for two weeks with the Cdk5i peptide resulted in significantly reduced levels of pTau compared organoids treated with the scrambled peptide (**Figure 4D-F**). Thus, the Cdk5i peptide counteracts pathological Tau hyperphosphorylation in human FTD models as it does in our transgenic mouse and cell-based systems.

## Discussion

We previously showed that genetic modification of p35 to abolish calpain-dependent cleavage and production of p25 successfully attenuated various pathological features associated with AD and FTD in their respective mouse models and cerebral organoids generated from patient-derived iPSCs (Seo et al., 2014; 2017). Non-cleavable mutant p35 (Δp35KI) is a very useful tool to study the role of Cdk5/p25 in various conditions and to elucidate mechanisms of pathogenesis for Cdk5-associated diseases, however, its application as a therapeutic is limited due to the technical and ethical challenges of applying genetic engineering technologies to human patients (Li et al., 2020). Here, we report the generation and initial characterization of a promising alternative treatment option.

Peptides are highly selective and relatively safe agents for therapeutic intervention. We designed a 12-amino-acid residue peptide based on our previous structural characterization of the Cdk5/p25 complex (Tarricone et al., 2001). We found that this peptide, termed Cdk5i, bound more than 40-fold more strongly to the Cdk5/p25 complex in *in vitro* assays than it did to either recombinant Cdk5 or Cdk2, suggesting it is highly specific for Cdk5/p25. Our findings also demonstrated that Cdk5i is capable of inhibiting Cdk5/p25 activity *in vitro* and Cdk5 activity *in vivo.* Importantly, this effect on Cdk5 activity *in vivo* was only observed on Cdk5 complexes isolated from Tau P301S brain lysates but not brain lysates from wild-type mice where Cdk5/p25 levels are expected to be very low. Further, consistent with the weak affinity Cdk5i showed for Cdk2 in our *in vitro* binding assay, we found that Cdk5i had no effect on the kinase activity of Cdk2 isolated from wild-type mouse brains. Thus, Cdk5i appears to show greater specificity for its intended target than existing small molecule inhibitors such as roscovitine, while also being considerably smaller than the peptide inhibitors of Cdk5 previously reported (Cicenas et al., 2015; Zheng et al., 2005; 2010).

We then explored the *in vivo* effects of Cdk5i by addition of a cell penetrating TAT sequence, as well as a FITC tag to monitor Cdk5i brain penetration and localization. Twenty-four hours after a single IP injection we were able to detect FITC-Cdk5i in both cortical and hippocampal sections, indicating the ability of this peptide to cross the blood-brain barrier to target Cdk5 complexes in the brain. Critically, we also observed striking effects of Cdk5i treatment on multiple early pathological phenotypes in CK-p25 mice, as well as on Tau phosphorylation in FTD-organoids, two neurodegenerative disease models characterized by elevated p25 production and Cdk5/p25 complex activity. Importantly, in addition to ameliorating elevated neuronal DNA damage in CK-p25 mice, Cdk5i treatment also reduced astrogliosis and microgliosis in CK-p25 brains, indicating at least partial reversal of the cascade of cellular changes that envelops all brain cell types during the course of neurodegeneration.

While we see clear effects of Cdk5i on Cdk5 complex activity and phenotypes *in vitro* and *in vivo*, its precise mode of action remains uncertain. Consisting of a stretch of Cdk5 residues predicted to mediate the interaction with p25, we hypothesized Cdk5i would bind primarily to p25 (and potentially also to p35). We indeed observed highly selective binding of Cdk5i to Cdk5/p25 complexes in our *in vitro* binding assay. We also observed pull-down of p35 by Cdk5i from wild-type mouse brain lysates, but intriguingly also Cdk5 itself, suggesting Cdk5i may bind independently to both Cdk5 and its activator proteins. While direct Cdk5i interactions with Cdk5 or its activators may serve to block Cdk5 complex formation, it is important to note that we observed that Cdk5i interacts more than 90-fold more strongly to Cdk5/25 complex than Cdk5 alone. Thus, it is possible that complexes containing all of Cdk5, activator and Cdk5i may form with impaired kinase function as opposed to the formation of Cdk5-Cdk5i, p25-Cdk5i or p35-Cdk5i complexes. Additional modes of action for Cdk5i are also conceivable and further experiments will be required to conclusively demonstrate the precise actions of this peptide. Although we saw preventative effects of our modified Cdk5i peptide *in vivo*, it is likely these functions can be improved further by identifying the key residues important for its inhibitory functions and making the peptide even smaller. Furthermore, adding auxiliary sequences may allow for effective delivery to target tissues/organs, increased stability, and prevent unwanted targeting by enzyme modifiers (Kaspar and Reichert, 2013; Fosgerau and Hoffmann, 2015).

The Cdk5i peptide efficiently inhibits Cdk5/p25 activity without affecting basal activity of Cdk5 important for neurodevelopment and physiological function of brain. Therefore, Cdk5i has potential to be applied to not only neurodegenerative diseases such as AD, PD, FTD mentioned earlier, but also various diseases associated with aberrant Cdk5 activity including cancer, stroke and neuropathic pain (Zhang et al., 2012; Meyer et al., 2014; Pozo and Bibb, 2016).

## Acknowledgements

We thank E. McNamara and M. Taylor for mouse colony maintenance and members of Tsai lab for discussion and valuable comments on manuscript. We also thank Harvard’s Center for Macromolecular Interactions for access to instrumentation for Biolayer Interferometry (BLI) and Microscale Thermophoresis. S.J.H. received funding from the Tau Consortium, Stuart & Suzanne Steele MGH Research Scholars Program, and Souvien Therapeutics. This work was supported by NIH grants NS051874-22, AG054012, NS102730, and Glenn award for research in biological mechanism of aging to L.-H.T.

## Figure Legends

**Supplementary Figure 1.** Demonstration of direct binding of the Cdk5i peptide to biotinylated Cdk5/p25 via biolayer interferometry. Top, the processed data graph with cdk5i as the analyte (n=5) following reference (n=3) subtraction show the summary of the real-time data acquisition in a binding kinetics experiment. After an initial baseline step in the assay buffer, streptavidin biosensors were dipped into solution with the biotinylated Cdk5/p25 for the loading of the protein onto the biosensors. Subsequently, a second baseline was performed, followed by association and dissociation of the analyte (n=5) in solution. A productive binding interaction leads to distinct spectral shift pattern that is represented on the sensorgram as a change in the wavelength (nm shift). The data were fitted globally with the 1:1 binding model and a representative plot of Req vs. Cdk5i concentration for the estimation of the equilibrium dissociation constant K_D_ is shown (bottom). Additional details on the BLI assay with biotinylated Cdk5/p25 were described previously (Patnaik et al., 2020).

**Supplementary Figure 2.** Schematics of isogenic control line generation from an FTD patient iPSCs carrying the *MAPT* P301L mutation. Sanger sequencing confirms the insertion of a desired repair template into an FTD patient iPSCs.

## References

Chae T, Kwon YT, Bronson R, Dikkes P, Li E, Tsai LH (1997) Mice lacking p35, a neuronal specific activator of Cdk5, display cortical lamination defects, seizures, and adult lethality. Neuron 18:29–42.

Cicenas J, Kalyan K, Sorokinas A, Stankunas E, Levy J, Meskinyte I, Stankevicius V, Kaupinis A, Valius M (2015) Roscovitine in cancer and other diseases. Ann Transl Med 3:135.

Fosgerau K, Hoffmann T (2015) Peptide therapeutics: current status and future directions. Drug Discov Today 20:122–128.

Gilmore EC, Ohshima T, Goffinet AM, Kulkarni AB, Herrup K (1998) Cyclin-dependent kinase 5-deficient mice demonstrate novel developmental arrest in cerebral cortex. J Neurosci 18:6370–6377.

Hall MD, Yasgar A, Peryea T, Braisted JC, Jadhav A, Simeonov A, Coussens NP (2016) Fluorescence polarization assays in high-throughput screening and drug discovery: a review. Methods Appl Fluoresc 4:022001.

Holmes BB, Furman JL, Mahan TE, Yamasaki TR, Mirbaha H, Eades WC, Belaygorod L, Cairns NJ, Holtzman DM, Diamond MI (2014) Proteopathic tau seeding predicts tauopathy in vivo. PNAS 111:E4376–E4385.

Kaspar AA, Reichert JM (2013) Future directions for peptide therapeutics development. Drug Discov Today 18:807–817.

Kim D, Frank CL, Dobbin MM, Tsunemoto RK, Tu W, Peng PL, Guan J-S, Lee B-H, Moy LY, Giusti P, Broodie N, Mazitschek R, Delalle I, Haggarty SJ, Neve RL, Lu Y, Tsai L-H (2008) Deregulation of HDAC1 by p25/Cdk5 in neurotoxicity. Neuron 60:803–817.

Kim SH, Ryan TA (2010) CDK5 serves as a major control point in neurotransmitter release. Neuron 67:797–809.

Kimura T, Ishiguro K, Hisanaga S-I (2014) Physiological and pathological phosphorylation of tau by Cdk5. Front Mol Neurosci 7:65.

Kolarova M, García-Sierra F, Bartos A, Ricny J, Ripova D (2012) Structure and pathology of tau protein in Alzheimer disease. Int J Alzheimers Dis 2012:731526.

Kusakawa G, Saito T, Onuki R, Ishiguro K, Kishimoto T, Hisanaga S (2000) Calpain-dependent proteolytic cleavage of the p35 cyclin-dependent kinase 5 activator to p25. J Biol Chem 275:17166–17172.

Lee MS, Kwon YT, Li M, Peng J, Friedlander RM, Tsai LH (2000) Neurotoxicity induces cleavage of p35 to p25 by calpain. Nature 405:360–364.

Li H, Yang Y, Hong W, Huang M, Wu M, Zhao X (2020) Applications of genome editing technology in the targeted therapy of human diseases: mechanisms, advances and prospects. Signal Transduct Target Ther 5:1–23.

Mazanetz MP, Fischer PM (2007) Untangling tau hyperphosphorylation in drug design for neurodegenerative diseases. Nat Rev Drug Discov 6:464–479.

Meyer DA, Torres-Altoro MI, Tan Z, Tozzi A, Di Filippo M, DiNapoli V, Plattner F, Kansy JW, Benkovic SA, Huber JD, Miller DB, Greengard P, Calabresi P, Rosen CL, Bibb JA (2014) Ischemic stroke injury is mediated by aberrant Cdk5. J Neurosci 34:8259–8267.

Moerke NJ (2009) Fluorescence Polarization (FP) Assays for Monitoring Peptide-Protein or Nucleic Acid-Protein Binding. Curr Protoc Chem Biol 1:1–15.

Ohshima T, Ward JM, Huh CG, Longenecker G, Veeranna Pant HC, Brady RO, Martin LJ, Kulkarni AB (1996) Targeted disruption of the cyclin-dependent kinase 5 gene results in abnormal corticogenesis, neuronal pathology and perinatal death. Proc Natl Acad Sci USA 93:11173–11178.

Park J et al. (2019) Abnormal Mitochondria in a Non-human Primate Model of MPTP-induced Parkinson’s Disease: Drp1 and CDK5/p25 Signaling. Exp Neurobiol 28:414–424.

Patnaik D, Pao P-C, Zhao W-N, Silva MC, Hylton NK, Chindavong PS, Pan L, Tsai L-H, Haggarty SJ (2020) Exifone is a Potent HDAC1 Activator with Neuroprotective Activity in Human Neuronal Models of Neurodegeneration. bioRxiv 2020:2020.2003.2002.973636.

Patrick GN, Zukerberg L, Nikolic M, la Monte de S, Dikkes P, Tsai LH (1999) Conversion of p35 to p25 deregulates Cdk5 activity and promotes neurodegeneration. Nature 402:615–622.

Pozo K, Bibb JA (2016) The Emerging Role of Cdk5 in Cancer. Trends Cancer 10:606–618.

Raja WK, Mungenast AE, Lin Y-T, Ko T, Abdurrob F, Seo J, Tsai L-H (2016) Self-Organizing 3D Human Neural Tissue Derived from Induced Pluripotent Stem Cells Recapitulate Alzheimer’s Disease Phenotypes. PLoS ONE 11:e0161969.

Ran FA, Hsu PD, Wright J, Agarwala V, Scott DA, Zhang F (2013) Genome engineering using the CRISPR-Cas9 system. Nat Protoc 8:2281–2308.

Seo J, Giusti-Rodríguez P, Zhou Y, Rudenko A, Cho S, Ota KT, Park C, Patzke H, Madabhushi R, Pan L, Mungenast AE, Guan J-S, Delalle I, Tsai L-H (2014) Activity-dependent p25 generation regulates synaptic plasticity and Aβ-induced cognitive impairment. Cell 157:486–498.

Seo J, Kritskiy O, Watson LA, Barker SJ, Dey D, Raja WK, Lin Y-T, Ko T, Cho S, Penney J, Silva MC, Sheridan SD, Lucente D, Gusella JF, Dickerson BC, Haggarty SJ, Tsai L-H (2017) Inhibition of p25/Cdk5 Attenuates Tauopathy in Mouse and iPSC Models of Frontotemporal Dementia. J Neurosci 37:9917–9924.

Shukla V, Seo J, Binukumar BK, Amin ND, Reddy P, Grant P, Kuntz S, Kesavapany S, Steiner J, Mishra SK, Tsai L-H, Pant HC (2017) TFP5, a Peptide Inhibitor of Aberrant and Hyperactive Cdk5/p25, Attenuates Pathological Phenotypes and Restores Synaptic Function in CK-p25Tg Mice. J Alzheimers Dis 56:335–349.

Shukla V, Zheng YL, Mishra SK, Amin ND, Steiner J, Grant P, Kesavapany S, Pant HC (2013) A truncated peptide from p35, a Cdk5 activator, prevents Alzheimer’s disease phenotypes in model mice. FASEB J 27:174–186.

Smith PD, Crocker SJ, Jackson-Lewis V, Jordan-Sciutto KL, Hayley S, Mount MP, O’Hare MJ, Callaghan S, Slack RS, Przedborski S, Anisman H, Park DS (2003) Cyclin-dependent kinase 5 is a mediator of dopaminergic neuron loss in a mouse model of Parkinson’s disease. Proc Natl Acad Sci USA 100:13650–13655.

Su SC, Tsai L-H (2011) Cyclin-dependent kinases in brain development and disease. Annu Rev Cell Dev Biol 27:465–491.

Tarricone C, Dhavan R, Peng J, Areces LB, Tsai LH, Musacchio A (2001) Structure and regulation of the CDK5-p25(nck5a) complex. Mol Cell 8:657–669.

Wei S et al. (2015) Active Pin1 is a key target of all-trans retinoic acid in acute promyelocytic leukemia and breast cancer. Nat Med 21:457–466.

Yoshiyama Y, Higuchi M, Zhang B, Huang S-M, Iwata N, Saido TC, Maeda J, Suhara T, Trojanowski JQ, Lee VM-Y (2007) Synapse loss and microglial activation precede tangles in a P301S tauopathy mouse model. Neuron 53:337–351.

Zhang HH, Zhang XQ, Wang WY, Xue QS, Lu H, Huang JL, Gui T, Yu BW (2012) Increased synaptophysin is involved in inflammation-induced heat hyperalgesia mediated by cyclin-dependent kinase 5 in rats. PLoS ONE 7:e46666

Zheng Y-L, Amin ND, Hu Y-F, Rudrabhatla P, Shukla V, Kanungo J, Kesavapany S, Grant P, Albers W, Pant HC (2010) A 24-residue peptide (p5), derived from p35, the Cdk5 neuronal activator, specifically inhibits Cdk5-p25 hyperactivity and tau hyperphosphorylation. J Biol Chem 285:34202–34212.

Zheng Y-L, Kesavapany S, Gravell M, Hamilton RS, Schubert M, Amin N, Albers W, Grant P, Pant HC (2005) A Cdk5 inhibitory peptide reduces tau hyperphosphorylation and apoptosis in neurons. EMBO J 24:209–220.

